# DISSECT: an assignment-free Bayesian discovery method for species delimitation under the multispecies coalescent

**DOI:** 10.1101/003178

**Authors:** Graham Jones, Bengt Oxelman

## Abstract

**Motivation:** The multispecies coalescent model provides a formal framework for the assignment of individual organisms to species, where the species are modeled as the branches of the species tree. None of the available approaches so far have simultaneously co-estimated all the relevant parameters in the model, without restricting the parameter space by requiring a guide tree and/or prior assignment of individuals to clusters or species.

**Results:** We present DISSECT, which explores the full space of possible clusterings of individuals and species tree topologies in a Bayesian framework. It uses an approximation to avoid the need for reversible-jump MCMC, in the form of a prior that is a modification of the birth-death prior for the species tree. It incorporates a spike near zero in the density for node heights. The model has two extra parameters: one controls the degree of approximation, and the second controls the prior distribution on the numbers of species. It is implemented as part of BEAST and requires only a few changes from a standard *BEAST analysis. The method is evaluated on simulated data and demonstrated on an empirical data set. The method is shown to be insensitive to the degree of approximation, but quite sensitive to the second parameter, suggesting that large numbers of sequences are needed to draw firm conclusions.

**Availability:** http://code.google.com/p/beast-mcmc/, http://www.indriid.com/dissectinbeast.html

**Contact:**, http://www.indriid.com

**Supplementary information:** Supplementary material is available.

## 1 Introduction

Despite its alleged status as a fundamental concept in biology, the species category has lacked a definition allowing explicit testing of particular species limits (e.g., de Queiroz, 2007). In recent years however, several methods have been proposed for the task of delimiting species based on molecular data (see Fujita *et al.*, 2012; Miralles and Vences, 2013, for reviews). Multispecies coalescent (Yang and Rannala, 2003) species limitation methods (MSCSLM) make use of multi-locus sequence data to make inferences in the presence of incomplete lineage sorting. We do not consider methods that only take the current genetic structure into account (e.g. STRUCTURE, STRUCTURAMA, Huelsenbeck and Andolfatto, 2007).

***Motivation:*** Ence and Carstens (2011) introduced the validatation/discovery terminology for species delimitation. All current MCSSLM are either heuristic (e.g. O’Meara, 2010), dependent on a guide tree (e.g. Yang and Rannala, 2010; Satler *et al.*, 2013) or are validation methods which require prior assignment of individuals to clusters or species. Here, we take a Bayesian approach, which has the advantage that nuisance parameters can be integrated out, and also that prior taxonomic knowledge can properly be taken to account.

***Our contribution:*** We present a Bayesian method DISSECT (Division of Individuals into Species using Sequences and Epsilon-Collapsed Trees) for species delimitation which requires no prior assignment of individuals to clusters or species, but instead explores the full space of possible clusterings and tree topologies. It is along the lines of the method of Yang and Rannala (2010) which employs a user-supplied guide tree in which some nodes may be collapsed (i.e., all descendants of these nodes assigned to one species). The two operations of collapsing a node, and of setting its height to zero, have the same effect on the likelihood, since the multispecies coalescent density is the same for a single population and a population which has just split at time zero. When a node is collapsed, the dimensionality of the parameter space changes, so a reversible-jump Markov Chain Monte Carlo (rjMCMC) algorithm is needed to sample the species trees. The basic idea behind DISSECT is to sample trees in which each tip represents a single individual (or a cluster of individuals which definitely belong in one species), but replace the usual prior density on node heights with one which includes a spike near zero. The dimensionality of the parameter space is fixed, but nodes whose heights have a high posterior probability of being within the spike can be interpreted as ‘probably collapsed’.

***Related work:*** Knowles and Carstens (2007) devised a Maximum Likelihood approach which used fixed gene trees as input data and hierarchical likelihood ratio tests to compare different species classifications. These were treated as different stochastic models with different sets of parameters, and the hierarchical likelihood ratio tests require the models to be nested. Thus, for example, the classification of putative species A, B, and C into AB and C or A and BC can not be compared in this way, whereas ABC can be compared to either. O’Meara (2010) devised parametric and non-parametric heuristic discovery methods to simultaneously find an optimal assignment of individuals to species and their tree relationships. Yang and Rannala (2010, 2013) developed the idea in a Bayesian framework, in which the gene trees are co-estimated with a constrained species tree, and implemented this in the software BP&P.

Two options are described in Yang and Rannala(2010). In the simplest option, the dimensionality of the parameter space does not change, and there is no special prior involved. Species are inferred by setting a threshold on the posterior node heights, with small heights interpreted as evidence for collapsing a node. This is similar to using *BEAST (Heled and Drummond, 2010) with each individual in its own ‘species’ in the XML file, and estimating the actual species afterwards. Yang and Rannala’s other (main) method employs a user-supplied guide tree in which some nodes may be collapsed. This drastically limits the number of possible species delimitations that are considered. DISSECT might loosely be described as ‘between’ or ‘combining’ these two options.

An alternative is to use Bayes Factors, which can be achieved from accurate marginal likelihood estimates (Xie *et al.*, 2011; Baele *et al.*, 2012). Grummer *et al.* (2013) and Aydin and Oxelman (submitted) used this approach to choose among species classifications, and Leaché *et al.* (2013) extended the approach to be used for SNP data.

## 2 Methods

A set of individual organisms will be called a **cluster**. Each possible cluster of individuals in the analysis is a candidate for constituting a species. A set of clusters which do not overlap one another and which together include all the individuals in the analysis will be referred to as a **clustering**. In an analysis using DISSECT, some sets of individuals may be grouped by the user as **minimal clusters**: these may be merged but never split. We use ‘gene’ in a loose sense, to mean an alignment of a sequence region which is assumed to be homologous and unlinked to other such regions. A ‘gene copy’ is a single row from such an alignment.

### 2.1 The model

In Bayesian phylogenetic analysis, a prior distribution over species trees is needed, and for rooted trees as used here, the reconstructed birth-death process (Gernhard, 2008) is most often used. It includes the Yule process as a special case. The process is assumed to begin at some time *t* in the past with a single species, and is conditioned on producing the observed number of species at present. The time *t* is called the **origin time** or **origin height**. Theorem 2.5 of Gernhard (2008), following Thompson (1975) shows that, conditioned on t, the speciation rate λ, and the extinction rate μ the density of the unordered node heights are independently and identically distributed (i.i.d.) and are also independent of the number of tips *k.* This nice mathematical property makes the present model tractable. Let the density of a node height *s* be *f* (*s*|*k*, *t*, λ, *μ*) = *f* (*s*|*t*, λ, *μ*). In the present model, *f* (*s*) is replaced with with a mixture of *f* (*s*) and another density *m*(*s*) for *s*:

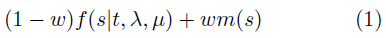

where *w* is a user-chosen weight in [0,1], and this density is used for all the *n* − 1 node heights in a tree with *n* tips. The joint density is then

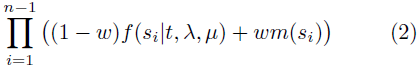
where *s*_1_, …, *s*_*n* − 1_ are unordered node heights. This can be expanded as

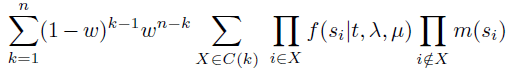

where *C(k)* is the set of subsets of {1,…, *n* − 1} of size *k* − 1. If *m*(*s*) was the Dirac delta function *δ(s)* (Dirac, 1958) the result would be a distribution in which the trees with *k* external branches of nonzero length (that is, the trees with *k* ‘real’ tips) have total probability mass

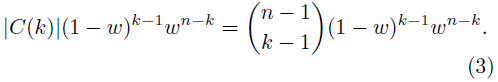

Note that the product 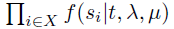 is the density for a reconstructed birth-death process with k tips whose node heights are the *k* – 1 nonzero s_i_. In practice one cannot sample from such a distribution without implementing reversible jump MCMC, but it can be approximated it using

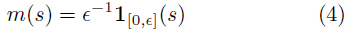
where ϵ is small.

Figures 1 and 2 illustrate the densities *f* and (1 – *w)f+wm* respectively for the case *n* = 3, where there are two internal node heights. One way of sampling trees from the reconstructed birth-death process for *n* = 3, is to pick a point (*x, y*) from a density such as the one in Figure 1; then choose a random ordering of the tip labels from left to right; then insert *x* and *y* between them; and finally join the nodes to form the tree. The same process is shown in Figure 2 for the mixture density *m*. If the point (*x, y*) is in one of the two ‘walls’ along the axes, one node will be collapsed. If the point (*x, y*) is in the ‘pillar’ near the origin, both nodes will be collapsed. The approximation means that there is a possibility that a true speciation which is more recent than ϵ, will be missed.

**Figure. 1.**
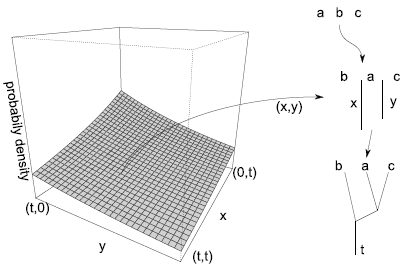
Sampling trees from the usual birth-death density.

**Figure. 2.**
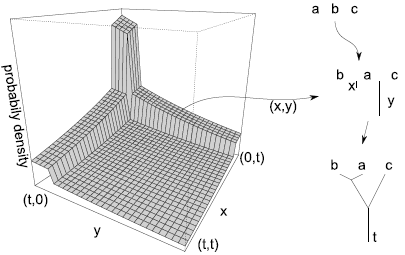
Sampling trees from the mixture density.

This is very similar to a model in which a separate reconstructed birth-death process is assumed for each *k* and a rjMCMC is used to sample from the clusterings and trees. Apart from the approximation involving ϵ, the other difference is that the density *q*(*t*|*k*) for *t* would normally depend on *k* in the reversible-jump version, whereas in equation (2) there is no such dependence: a single density for *t* for all *k* is needed. It seems reasonable to assume a density for *q(t)* which mixes *q*(*t*|*k*) using the probabilities from expression (3). In a normal BEAST or *BEAST analysis using the birth-death prior, an improper uniform prior on [0, ∞) is assumed for the origin time *t* of the tree, and the process is then conditioned on the number of species *k*. The conditional density for *t* is shown in Theorem 3.2 of Gernhard (2008) to be

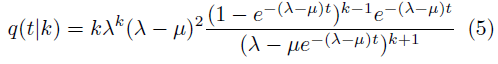

Using the probabilities from expression (3), the prior density for *t* is

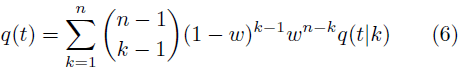

This can be simplified as shown in the supplementary information.

The model was implemented in BEAST by adding a class BirthDeathCollapseModel, which is similar to the usual BirthDeathModel. It contains a parameter for the origin height *t* as well as for the diversification rate and relative death rate as in the usual birth-death model. An additional MCMC operator is needed to sample from *t*. This can be added using one of the existing operators in the XML. We used a ScaleOperator. No new MCMC operators were added to explore the space of species trees: the existing NodeReHeight operator explores the posterior as modified by the prior in equation (equation 2).

### 2.2 DISSECT workflow

The analysis can be run in the development version of BEAST (Drummond *et al.*, 2012), see supplementary data for instructions. BEAUTi can be used to set up most of the analysis, as if for a *BEAST analysis. The word ‘species’, as it appears in BEAUTi and in the BEAST XML file, is interpreted as a minimal cluster. Two changes need to be made to the XML file. The birth-death model must be replaced with a birth-death-collapse model, where ϵ can be set, and an operator must be added for the origin height. The parameter *w* can either be given a fixed value, or estimated by adding a hyperprior and an operator. The trees sampled from the posterior can be analyzed with a tool called SpeciesDelimitationAnalyser. This uses a user-supplied threshold ϵ (either ϵ or larger) for assigning individuals to clusters, and produces a table *H* of clusterings *Z*_1_, *Z*_2_,… *Z*_z_ with corresponding posterior probabilities *p*_1_, *p*_2_,…*p*_z_ which sum to 1. The clusterings are sorted in order of decreasing posterior probability. An R script for producing a similarity matrix (see section 3.4) is provided in the supplementary information.

### 2.3 Advice on choosing parameters and priors

The parameter *w* can be chosen to reflect prior knowledge about the likely number of species. As a consequence of the structure of the model, even when *w* is fixed, the prior on the number of species *k* is somewhat diffuse: it is not possible to insist on exactly 7 species for example. In the case of fixed *w*, the number of trees with *k* ‘real’ tips in the prior has the distribution of 1+*X* where *X* is a random variable having the binomial distribution with size parameter *n* − 1 and probability parameter 1 − *w*. Its mean is thus 1+(*n* − 1)(1 − *w*). If the individuals have been assigned in previous work to *k_0_* species, then *w* = (*n* − *k*_0_)/(*n* − 1) seems a reasonable choice. If the value of *w* is estimated, and a beta prior is used, the prior distribution on *k* − 1 is a beta-binomial distribution, which can be explored using the R package VGAM (Yee, 2010,, see also supplementary information). If *w* is fixed at zero, the value of *ϵ* becomes irrelevant, and the model becomes equivalent to the birth-death model as used in *BEAST, except that the origin height is estimated instead of being integrated out analytically.

The parameter ϵ should be set to a small value such as 1e-4 or 1e-5. The value is a compromise between exactly matching a particular model and the practicalities of computation. Extremely small val-ues may lead to poor mixing, although we have only observed a substantial effect for ϵ below 1e-6. If ϵ is too large it will not be possible to distinguish very recent divergences. For most analyses there will not be enough data to distinguish speciations with node heights below 1e–4, since the expected number of mutations separating the species is only one per 5,000 sites, so the choice of ϵ will not be at all critical.

When the number of individuals per species is small, it becomes difficult to estimate the population size parameters in each branch in the multispecies coalescent model. In such a case, care must be taken to use a sensible prior on these parameters, especially the ‘species.popMean’ parameter. We recommend that the prior should be proper, and diffuse enough to accommodate extreme but possible values, but not absurdly diffuse. This is good advice anyway when using *BEAST, but it becomes more critical when using DISSECT, since it will typically be harder to ensure that the number of individuals per species is not small.

## 3 Evaluation

### 3.1 Simulated scenarios and parameter settings

Two sets of simulations were run. The first set evaluates the performance of DISSECT as the number of genes and the amount of incomplete lineage sorting varies, and assesses the sensitivity of the method to choices of ϵ and *w*. The second set focuses on the case of one true species. We use *N_e_* to mean the effective number of (diploid) individuals in a population. If *N_e_* is constant, this means that the expected time for two gene copies to coalesce is *2N_e_* generations. We denote the mutation rate per site per generation by *T*. Node heights and ϵ are in the same units as the product *TN_e_.* Note that the topology and node heights of the gene trees only depend on the product *TN_e_*, so a scenario with *T* = 1e-8 and *N_e_* = 50,000 is equivalent to one with *T* = 1e-9 and *N_e_* = 500,000 and so on. There are two sources of ‘noise’ in the data: one comes from coalescences which are deeper than species tree node height, and the other from the randomness of mutations. For a node height of 0.001 and a gene length of 500, the expected number of substitutions separating two species is 1, so around 37% of pairs of gene copies from different species would be identical if they coalesced at the species node height. In all cases the threshold ϵ in SpeciesDelimitation-Analyser was set equal to ϵ

The first set SIM-5×5 of simulations all use 25 individuals, 5 assigned to each of 5 species, with one gene copy per individual. The species tree has a comb topology with node heights at 0.001, 0.002, 0.004, and 0.008. These heights are chosen to roughly approximate those in the empirical data set (see below). The value of *N_e_* was 50,000 at the tips and at the root-ward ends of branches, and 100,000 at the root and tipwards ends of internal branches, varying linearly along the branches. The length of the genes was set to 500 sites, and the number of genes *G* was set to 3 or 9. The mutation rate *T* was set to 1e-8 or 4e-8, representing a moderate and a large amount of incomplete lineage sorting. The root of the species tree is at 0.008/T generations, and is therefore 800,000 generations when *T* = 1e-8 and 200,000 generations when *T* = 4e-8. To get some idea of the amount of signal and noise due deep to coalescences in the data, consider the *G* = 9 case, where there are 4,500 sites. For a very small value of *T*, the number of variable sites would be about 100. In the case *T* = 1e-8, the number of variables site was around 200, and in the case *T* = 4e-8, it was around 400. The increase in variable sites as *T* increases is due to deeper coalescences.

We explored the accuracy of the method with respect to changes in ϵ by using a beta prior for *w* with shape parameters 8 and 2, and setting ϵ to 0.0001 = 1e-4, 3e-5, and 1e-5. We also explored the behavior with respect to changes in *w* by fixing ϵ to 1e-4, and setting *w* to 11/12, 5/6, and 17/24, corresponding to prior means for *k* of 3, 5, and 8.

The second set SIM-1 of simulations all use *T* = 1e-8 and *N_e_* = 100,000. In this case a single species was simulated, so the gene trees are all the result of a coalescence process only. The product *TN_e_* scales the number of substitutions, and thus affects the accuracy with which genes trees can be estimated, but does not change the underlying ‘shape’ of the problem. The value of ϵ was 3e-5. A beta prior with shape parameters *n* − 1 and 1 was used for *w*, which means that the true clustering has a probability of 0.5 in the prior for all *n*. We used *n* = 4, 8, and 16 individuals and *G* was set to 3 and 9 to examine how these variables affect the rate of false splits.

### 3.2 Implementation of simulations

The simulated data was generated and analysed using R (R Development Core Team, 2011) and the R packages APE (Paradis, 2004) and phangorn (Schliep, 2011). Gene trees were simulated according to the multispecies coalescent model for each scenario and parameter choice, for ten replicates. Sequence alignments with 500 sites were generated for these gene trees using Seq-Gen Rambaut and Grassly (1997) called with command seqgen.exe -`mHKY` -t3.0 -f0.3,0.2,0.2,0.3. This uses a strict clock and the HKY substitution model, and all genes have the same mutation rate. There is no site rate heterogeneity. These sequences were then incorporated into BEAST XML files, and DISSECT was run for 20 million generations with the first 10 million discarded as burnin. The priors for `species.popMean`, `meanGrowthRate`, and the relative clock rates were all lognormals, with means and standard deviations in log space equal to -7 and 2; 4.6 and 2; and 0 and 1 respectively. The prior for `relativeDeathRate` was uniform in [0,1]. SpeciesDelimitationAnalyser was run after DISSECT.

### 3.3 Empirical data

Species delimitation in the pocket gopher genus *Thomomys* subgenus *Megascapheus* has been controversial, with a large number of species described by early taxonomists. Most of these have been reduced to subspecific rank by 20th century taxonomists inspired by the biological species concept (e.g., Wilson and Reeder, 2005). According to opinion of these recent authors, the number of species in the dataset of Belfiore *et al.* (2008), also used by Heled and Drummond (2010), vary between six and eight, depending on how the species *T. bottae, T. umbrinus*, and *T. townsendii* are delimited. We explored the dataset, which consists of 26 individuals and seven non-coding nuclear sequence regions (Belfiore *et al.*, 2008), by varying ϵ from 1e-7 to 1e-3=0.001, and by setting *w* to 0.12 or 0.68 (corresponding to subspecies elevated to species rank, and eight species, respectively). We also used a Beta hyperprior with parameters 4 and 2 (Fig. 3). Each combination of ϵ and *w* was run for 200 million generations, saving parameter values and species trees every 5000th generation. SpeciesDelimationAnalyser was run with ϵ equal to ϵ, and one magnitude of order larger.

**Figure. 3.**
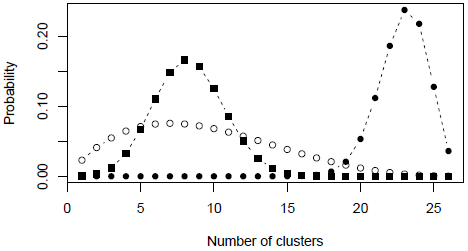
Prior distribution for the number of clusters when *w* is 0.12 (black circles), 0.68 (black squares), and has a Beta distribution with parameters 4 and 2 (open circles).

### 3.4 Evaluation metrics

The number of possible clusterings of *n* individuals (known as the Bell number *B*_n_) increases rapidly with *n*. For example *B*_2_ = 2, *B*_3_ = 5, *B*_4_ = 15, *B*_5_ = 52, *B*_10_ = 115975, and *B*_25_ ≈ 4.6e18. (See O’Meara, 2010, for more details.) The accuracy of the estimated number of species is not a good way to judge the method, since the number may be correct despite false splits and false merges which cancel out, or despite major misassignments, or incorrect due to a single individual being incorrectly merged or separated from a cluster. There are approximately 2.4e15 ways in which 25 individuals can be grouped into 5 clusters. The situation is similar to that of inferring phylogenies, where we typically do not expect every clade to be correctly inferred if the number of species is large. In order to assess the accuracy of DISSECT, we therefore want a metric analogous to tree metrics such as the Robinson-Foulds distance.

***Rand index.*** We chose the Rand index (Rand, 1971), which measures the similarity *R*(*X,Y*) between two clusterings *X* and *Y* of the same set (e.g., the set of individuals). The Rand index is always between 0 and 1, and is 1 when the match is perfect. We also define 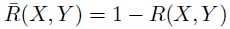 which is a metric in the mathematical sense, and which we will refer to as the Rand metric. Firstly, in order to evaluate the posterior distribution as a whole, we weight the Rand metric between each clustering *Z_m_* in the table Ω produced by DISSECT and the true clustering *Z\** by its posterior probability *p_m_*, and thus produce an overall measure of the distance from the posterior distribution to *Z*:*:

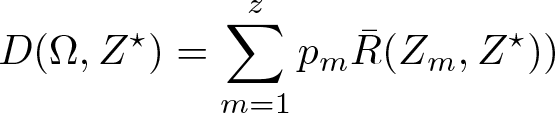

This is our main tool for evaluating DISSECT on simulated data.

***Point estimator.*** The simplest point estimator of the species limits, which we denote by *Z*, is the clustering with the highest posterior probability, that is, the posterior mode. An alternative point estimator is described in the supplementary information.

*Similarity matrix.* For any clustering *Z* in Ω, define mat(Z) to be the symmetric matrix which has (*i,j*)th entry equal to 1 if *i* and *j* are in the same cluster in *Z* and 0 otherwise. We define the similarity matrix to be the weighted sum of such matrices in Ω:

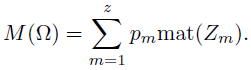

The elements of *M*(Ω) are the posterior probabilities for pairs of individuals to belong to the same cluster. This matrix can be thought of as a posterior mean in 'similarity matrix space’. It makes a convenient visual summary of the output from DISSECT. The point estimators and *M*(Ω) are the main tools for interpreting the output of DISSECT on empirical data.

***Credible sets.*** We define a distance between matrices *A* and *A* as the mean absolute difference of off-diagonal elements between the two matrices:

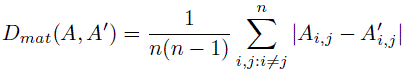

We calculate *D_mat_(Z, M*(Ω)) for each clustering *Z* in Ω and sort the clusterings on these distances in increasing order. Then we add clusterings to the credible set until the sum of posterior probabilities reaches the required size (e.g., 0.95). A single clustering may be partly in and partly outside the credible set in which case we regard it as belonging to the credible set if half its posterior probability lies within the required size. We chose this credible set in preference to a highest posterior density credible set because clusterings which only occur once in Ω can account for a large fraction of the posterior probability mass, and the distance from *M*(Ω) provides a sensible criterion for including or excluding them.

Finally we note some connections between these quantities. If *A* = mat(Z) and *A’* = mat(Z') for some clusterings *Z* and *Z'*, then *D_mat_(A, A')* is equal to 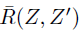. Denote the true similarity matrix mat(Z*) by *M*.* Then the distance between the true and estimated similarity matrices, namely *D_mat_(M*(Ω),M*) is equal to *D(Ω,Z\**) (as shown in the supplementary information) so *D*(Ω, *Z*\*) can be interpreted in two ways.

### 3.5 Results

Results for the first set SIM-5x5 are shown in Fig. 4 (varying ϵ) and Figs. 5, 6, 7, (varying *w*) and (table 1. These show no obvious differences when ϵ is changed over an order of magnitude. Other figures supporting this conclusion are in the supplementary information. The results for varying *w* show some effect, at least with the *G* = 9, *T* = 1e-8 case in Figs. 5 and 7.

**Table I.**
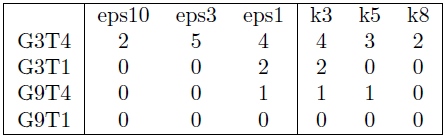
Number of times out of 10 that the true clus-tering failed to be in the 0.95 credible set, as G, T, ϵ and ϵ (first 3 columns) and *w* (last 3 columns) vary.

|  | eps10 | eps3 | eps1 | k3 | k5 | k8 |
| --- | --- | --- | --- | --- | --- | --- |
| G3T4 | 2 | 5 | 4 | 4 | 3 | 2 |
| G3T1 | 0 | 0 | 2 | 2 | 0 | 0 |
| G9T4 | 0 | 0 | 1 | 1 | 1 | 0 |
| G9T1 | 0 | 0 | 0 | 0 | 0 | 0 |

**Figure. 4.**
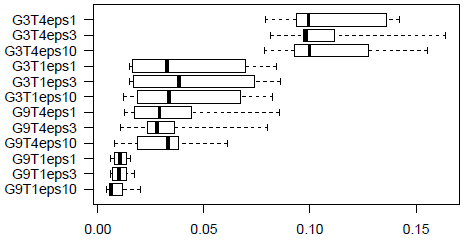
The boxplots show the values of the error metric *D*(Ω,*Z*\*.) over ten replicates as G, T, and ϵ. vary. In the labels, G3T4eps1 means the number of genes G is 3, the mutation rate T is 4e-8, and ϵ is 1e-5 = 0.00001. The other values of ϵ are 0.00003 and 0.0001. A beta prior with shape parameters 8 and 2 was used for *w* which is estimated.

**Figure. 5.**
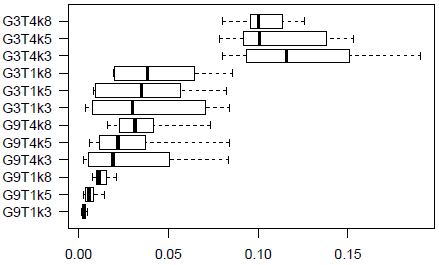
D(Ω,Z*.) as G, T, and *w* vary. The numbers after the ‘k’ in the label are the prior mean values for the number of species which are affected by the value of *w* The value of ϵ is 0.0001.

**Figure. 6.**
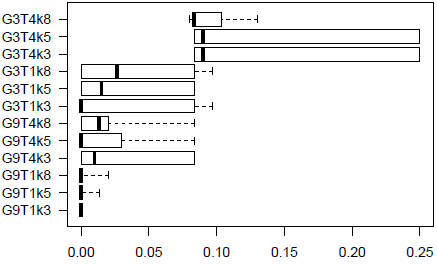
Rand metric between point estimate Z and true clustering, as G, T, and w vary. Other details as Fig. 5

**Figure. 7.**
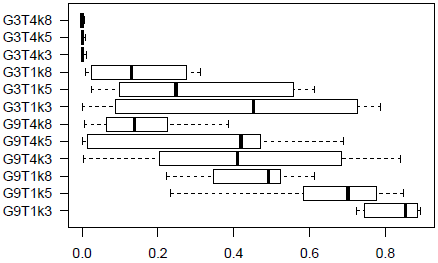
Posterior probability of true clustering, as G, T, and *w* vary. Other details as Fig. 5.

Results for the second set SIM-1 are shown in Figs. 8 and 9. The point estimates were always correct. There were no false splits with high posterior probabilities. The posterior probability of the correct clustering increases with both *n* and *G*.

Varying ɛ between 1e-4 and 1e-7 on the *Thomomys* data did not have any noticeable effects on the similarity matrices generated from SpeciesDelimitationAnalyzer (Fig. 10). Setting *τ* one magnitude of order higher than *ɛ* increased posteriors for clusterings in general, but not as much as increasing ϵ tenfold and setting *τ* equal to ϵ (supplementary information). Varying *w* had clear effects, with more and smaller clusters for the small w = 0.12 (Fig. 10). The posterior mean values for *w* when estimated with a Beta(4,2) prior distribution varied between 0.53 and 0.55 when ϵ was in the range not affecting the posteriors. Effective sample sizes (ESS) for most parameters were well above 300, except for some population size parameters for individual branches, and speciation.likelihood, where the smallest ϵ values gave low ESSs for w = 0.68 and *w* estimated with Beta(4,2). Fig. 11 shows the similarity matrix for ϵ = 1e-5 and *w* ~Beta(4,2).

**Figure. 8.**
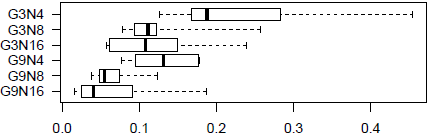
The true clustering is a single cluster. The plot shows D(Ω,Z*) for varying number of individuals n and genes G. The value of ϵ is 3e-5. A beta prior with shape parameters n-1 and 1 was used for *w* which is estimated.

**Figure. 9.**
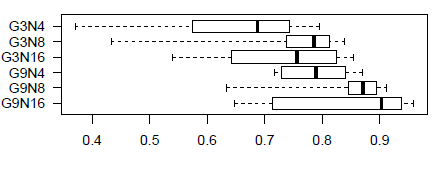
The true clustering is a single cluster. The boxplot shows the posterior probability of this, as *n* and G vary. Other details as Fig. 8.

**Figure. 10.**
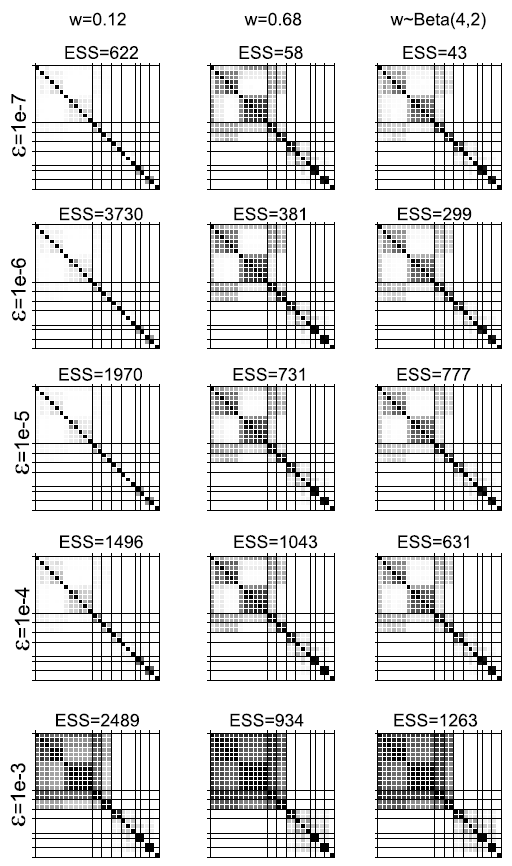
Similarity matrices for the Thomomys data set under various ϵ and collapse weight (*w*) values. The squares represent posterior probabilities (white = 0, black = 1) for pairs of individuals to belong to the same cluster. The ESS values are effective sample sizes for speciation.likelihood. The right column shows results when a Beta prior distribution with parameters 4 and 2 was used.

**Figure. 11.**
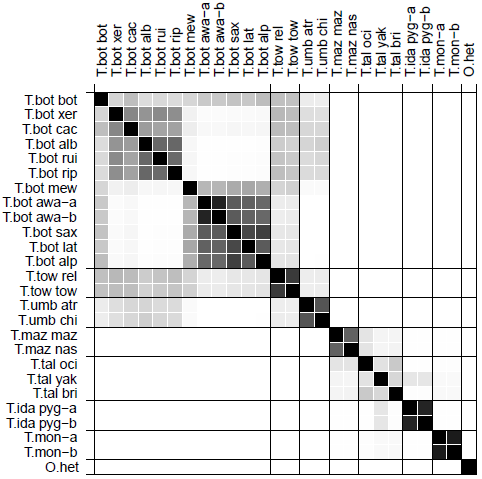
Similarity matrix for the Thomomys data set under ɛ=1e-5 and *w*~Beta(4,2). In the labels, ‘T’ stands for *Thomomys*, ‘O’ for Orthogeomys and species and subspecies designations are abbreviated to the first three letters.

## 4 Discussion

### 4.1 Simulations

As expected, the accuracy increases with the number of unlinked loci and the ability to detect species increases as the height of the nodes increases. The insensitivity of the method to varying ϵ suggests that the approximation is unlikely to bias the results.

The results in set SIM-5×5 for varying *w* in *G* = 9, *T* = 1e-8 case Fig 5, Fig 6, Fig 7) seem surprising, since this is the easiest case where the influence of the prior is expected to be least. One explanation is that with less data or more deep coalescences, there are a mixture of merge and split and mis-assignment errors, and changing *w* increases some and decreases others. When G = 9, *T* = 1e-8, merge errors are very rare, so a bias towards merging in the prior is reduces the overall error.

In table 1, the G=3,T=4e-8 results are poor, but this may be because there are a huge number of clusterings with tiny posterior probabilities, making it difficult to estimate the credible set. The true clustering may be in the true credible set, but not in the credible set that is estimated from ‘only’ 10,000 samples. We tried ten longer runs of 60 million generations, with 10 million discarded as burnin, and every 1000th generation sampled, for one of the *G* = 3,*T* = 4e-8 scenarios (eps3) and the true clustering was then found in the credible set in all ten runs.

The results on the scenarios SIM-1 with one true species show the method does not often infer false splits, but it is also clear that a substantial number of sequences are required in order to draw a firm conclusion even in this simplest of cases. A full evaluation of the method on more complex cases is beyond the scope of this paper. Note that even with two true species, there is a 4-dimensional space of scenarios to explore (node height, effective population size, number of individuals, and number of loci).

In general, the number of species was overestimated in the scenarios used here (results not shown). However, one could add very recent nodes to the scenarios which would tend to be falsely merged and result in an under-estimate instead. It would be interesting to evaluate the method on a large number of scenarios produced by sampling from a birth-death process. For the moment we suggest that estimates of numbers of species are treated with caution.

### 4.2 Empirical example

The insensitivity of DISSECT to ϵ suggested by the results from the simulated data seems corroborated by the Thomomys data. In data sets of the size evaluated here, there is far too little information to detect node heights smaller than 0.0001 substitutions per site. On the other hand, the impact of *w* on the data was noticeable, indicating that the data is not informative enough to be strongly conclusive about species delimitations. However, the possibility to include previous taxonomic opinions about species delimitations in the prior is promising, although it allows only species numbers, not species assignments to be considered.

The ambiguous assignment of several individuals in the *Thomomys* data set may indicate violations to the assumptions of the model (e.g., no hybridizations), or that the data is not informative enough. To assess absolute fit of the data to the model, posterior predictive simulation-based model checks may give clues to the reasons for this (Reid *et al.*, 2013). Indeed, the *Thomomys* data showed poor fit to the multispecies coalescent model in the survey by Reid *et al.* (2013), and one possible reason to this might be mis-assignment of alleles to species.

The multispecies coalescent model assumes no migration after speciation, which is instantaneous. This is probably violated in most cases. Zhang *et al.*(2010) found that low rates (< 0.1 migrant per generation) of migration had virtually no effect on the accuracy of BP&P in a two-species simulation study. However, at least when sample size is small, a single sampled recent migrant can cause severe effects. The coalescent prior on the gene trees will affect them in a way that single recent introgressions will be “pushed back” by other gene trees that reflect the “true” speciation event, such that the coalescent time for the migrant may be biased. More research is needed to evaluate the robustness of the model to hybridization, and in particular perhaps, to gradual isolation of species, which may be the most common form of speciation (e.g., Barton and Charlesworth, 1984). If this is true, it may be necessary to develop models that better fit such data.

### 4.3 Conclusion

‘Given the intrinsic theoretical and empirical difficulties of the problem, any success would be surprising.’ (O’Meara, 2010, p68). We believe that DISSECT is a useful step forward on the theoretical and computational side. The multispecies coalescent model has assumptions that are likely to be violated and it remains to be seen how important these are for empirical data.

We have not formally evaluated the accuracy of the species trees produced by DISSECT. However, apart from the approximation involving ϵ, and the slightly different prior on the tree root height, the DISSECT model, when conditioned on a particular clustering *Z*, is equivalent to *BEAST using *Z* to assign individuals to species. This means that DISSECT can be used as in a regular *BEAST analysis, taking uncertainties in species delimitation into account.

## Acknowledgement

We thank three anonymous reviewers for valuable comments on an earlier version of this paper.

